# Plasticity in parental behavior and vasopressin: Responses to co-parenting, pup age, and an acute stressor are experience-dependent

**DOI:** 10.1101/2023.03.01.530631

**Authors:** Lisa C. Hiura, Vanessa A. Lazaro, Alexander G. Ophir

## Abstract

The impact of variation in parental caregiving has lasting implications for the development of offspring. However, the ways in which parents impact each other in the context of caregiving is comparatively less understood, but can account for much of the variation observed in the postnatal environment. Prairie voles (*Microtus ochrogaster*) demonstrate a range of postnatal social groups, including biparental pairs and pups raised by their mothers alone. In addition to the challenges of providing parental care, prairie vole parents often experience acute natural stressors (e.g., predation, foraging demands, thermoregulation) that could alter the way co-parents interact. We investigated how variation in the experience of raising offspring impacts parental behavior and neurobiology by administering an acute handling stressor on prairie vole families of single mothers and biparental parents over the course of offspring postnatal development. Mothers and fathers exhibited robust behavioral plasticity in response to the age of their pups, but in sex-dependent ways. Pup-directed care from mothers did not vary as a function of their partner’s presence, but did covary with the number of hypothalamic vasopressin neurons in experience-dependent ways. The relationship between vasopressin neuron numbers and fathers’ behaviors was also contingent upon the stress handling manipulation, suggesting that brain-behavior associations exhibit stress-induced plasticity. These results demonstrate that the behavioral and neuroendocrine profiles of adults are sensitive to distinct and interacting experiences as a parent, and extend our knowledge of the neural mechanisms that may facilitate parental behavioral plasticity.

## 1 Introduction

Parents adapt their behaviors in response to environmental contexts, and one of the most influential contexts that a parent will experience is the dynamic social environment of familial interactions (Royle et al., 2014). All mammalian mothers interact with their offspring to varying degrees over the course of offspring development, but for the 5-10% of mammalian species that are biparental (Kleiman & Malcolm, 1981), partner interactions make up a substantial portion of the parents’ social environment. Several species (primarily studied in birds) have been found to flexibly respond to the actions or presence of a parenting partner (Harrison et al., 2009). Adapting parental behavior to meet the demands of a variable social environment can increase reproductive fitness, but social environmental variation can have long term consequences on the developing offspring and the parental investment they receive. Moreover, the dynamic responses of parents to the social environment have the potential to exhaust parental effort and impact brain function and behavioral phenotype. Yet, the ways in which caregiving and the maternal (and paternal) brain adapts to having a co-parent remain underexplored in mammals.

Prairie voles (*Microtus ochrogaster*) are small rodents that form socially monogamous pair bonds and exhibit biparental care. With the exception of nursing, fathers exhibit all of the same caregiving behaviors as mothers (Thomas & Birney, 1979). Nevertheless, paternal care is not necessary for pup survival in prairie voles, and the frequency that biparental pairs are observed in nature is about the same as the frequency that single mothers raise pups alone (Getz et al. 1993). Moreover, several studies have removed fathers from the natal nest to assess the consequences of variation in the early social environment on offspring development. These studies have determined that the presence of a father during the rearing period has significant impacts on pup development, including rates of physical maturation (Wang & Novak, 1992), alloparental care and pair bonding as adults (Ahern & Young, 2009), parental behavior toward their own offspring (Ahern et al., 2011), and neuropeptide receptor binding densities and mRNA expression (Ahern & Young, 2009; Bales & Saltzman, 2016; Prounis et al., 2015).

Interestingly, far fewer studies have investigated the role of father-presence on mammalian mothers. This is likely due to the comparatively fewer numbers of biparental species suitable for these inquiries. One such study in prairie voles found that single mothers show more passive stress-coping and anxiety-like behaviors alongside greater mRNA expression of hypothalamic corticotropin-releasing hormone compared to mothers who remained paired with their mate (Bosch et al., 2018). When maternal behaviors were compared in the days following parturition, no group differences were found. Other studies have found that even though male prairie voles are active participants in caregiving, mothers consistently provide high levels of care to offspring, whereas caregiving across fathers varies widely (Kelly et al. 2020; Finton & Ophir 2020; Solomon 1993). These findings reveal that variation in parental experience induced by the removal of the parenting partner has substantial context-specific behavioral and neuroendocrinological consequences for prairie vole mothers during the perinatal period.

Parental behaviors in male and female monogamous rodents are in part regulated by the neuromodulatory hormone vasopressin (VP; Horrell et al., 2018). Vasopressin is largely produced in the paraventricular nucleus (PVN) and the supraoptic nucleus (SON) of the hypothalamus, and these primary cell groups provide the bulk of VP peptide signals in the central and peripheral nervous systems (Brownstein et al., 1980). Interestingly, VP mRNA levels in the PVN and the SON are greater in prairie vole parents compared to sexually naïve controls (Wang et al., 2000). Moreover, when examining several brain regions implicated in the modulation of social behavior, prairie vole mothers and fathers show higher levels of OT-cFos and VP-cFos colocalization in the presence of pups compared to when they are separated from their litter or exposed to an object control, demonstrating that these cell populations are transcriptionally responsive to pup stimuli in both sexes (Kenkel et al. 2012; Kelly et al. 2017; Kirkpatrick, Kim, & Insel 1994). Critically, VP is not only involved in parental behavior; a variety of stressors induce the secretion of VP from the PVN, which functions as a key regulator of hypothalamic-pituitiary-adrenal axis responsivity (Herman & Tasker, 2016). For example, an acute swim stressor increased VP release within the PVN and SON of adult male rats (Wotjak et al. 1998) and heightened pup-directed care in male prairie voles (Bales et al., 2006). In addition, rats bred for high-anxiety traits had increased expression of VP mRNA in hypothalamic nuclei, and antagonism of PVN VP receptors reduced anxiety-like behaviors in such rats (Wigger et al., 2004). Together these findings suggest a functional role for VP in mediating endocrine and behavioral stress-responses, which may convolute our understanding of VP’s role in the simultaneous modulation of parental behaviors (Saltzman et al., 2017).

To better dissect the potential intersections between parental experience-dependent plasticity, stress, and the involvement of VP, we investigated how environmental variation during parenting impacts the behaviors of prairie vole mothers and fathers across the pre-wean stage of offspring development. Specifically, we implemented a social context manipulation (father removal) and an acute stress induction paradigm (experimenter handling) to produce diversity in parental rearing experiences. We chose experimenter handling because brief scruffing in rodents produces stress-induced catalepsy (Amir, Brown, Amit, & Ornstein 1981) and transient stress-related increases in heart rate and body temperature (Cinelli et al., 2006). Routine laboratory handling has also been found to induce elevations in the hormones corticosterone and prolactin, both of which are strongly implicated in physiological stress responses (Balcombe, Barnard, & Sandusky, 2004). Importantly for the present study, prairie voles that were directly handled by an experimenter showed elevated pup-directed care behaviors when compared to prairie voles that were simply transferred in cups (Tyler, Michel, Bales, & Carter, 2005). Here, we leverage this handling stress paradigm in conjunction with our manipulation of the presence or absence of fathers to diversify parental care experiences. We measured parents’ homecage behaviors and parent performance in an open field test, then subsequently immunolabeled VP-ir neurons in the PVN and SON to characterize the relationship(s) between biobehavioral plasticity and experiences as a parent.

## 2 Materials and Methods

### 2.1 Experimental Animals

All animals used in these experiments were sexually mature, virgin F1 progeny of wild-caught breeders and were reared in the laboratory at Cornell University. All animals had *ad libitum* access to water and food (Rodent Chow 5001, LabDiet, St. Louis, MO, USA) and were housed in polycarbonate rodent cages (29 × 18 × 13 cm) lined with Sani-chip bedding under a 14:10 light-dark cycle. Ambient temperature was maintained at 20C ± 2. Sex was assessed and assigned at weaning based on differences in external genitalia. All procedures were approved by the Institutional Animal Care and Use Committee of Cornell University (2013-0102) and were consistent with the guidelines set forth by the National Institutes of Health.

### 2.2. Family conditions

We created two experimentally designed factors to manipulate the animals: Fathers Present/Absent, and Experimentally Handled or Non-handled. Twenty days after breeding pairs were formed, males in families assigned to the Father-Absent condition were permanently removed from the home cage. Males from families assigned to the Father-Present condition were briefly removed, then returned to the home cage to control for nest disturbances. Nests were monitored daily for the birth of pups. At birth, litters were culled to three pups to control for the effect of litter size. Male offspring were preferentially spared for use in a separate related study. Weekly cage changes were conducted on postnatal days (PND) 2, 9, and 16. In the Handled condition, parents were scruffed and transferred by a gloved hand to a clean cage. In the Non-handled condition, families were gently scooped into a plastic beaker and transferred to a clean cage. When pups were not attached to their mothers, they were transferred in the same manner as their parent(s). Altogether we created four groups: Father Present / Non-handled (N = 11), Father Absent / Non-handled (N = 9), Father Present /Handled (N = 14), Father Absent / Handled (N = 8); see **Figure 1**. All pups were weaned at PND21.

**Figure 1.**
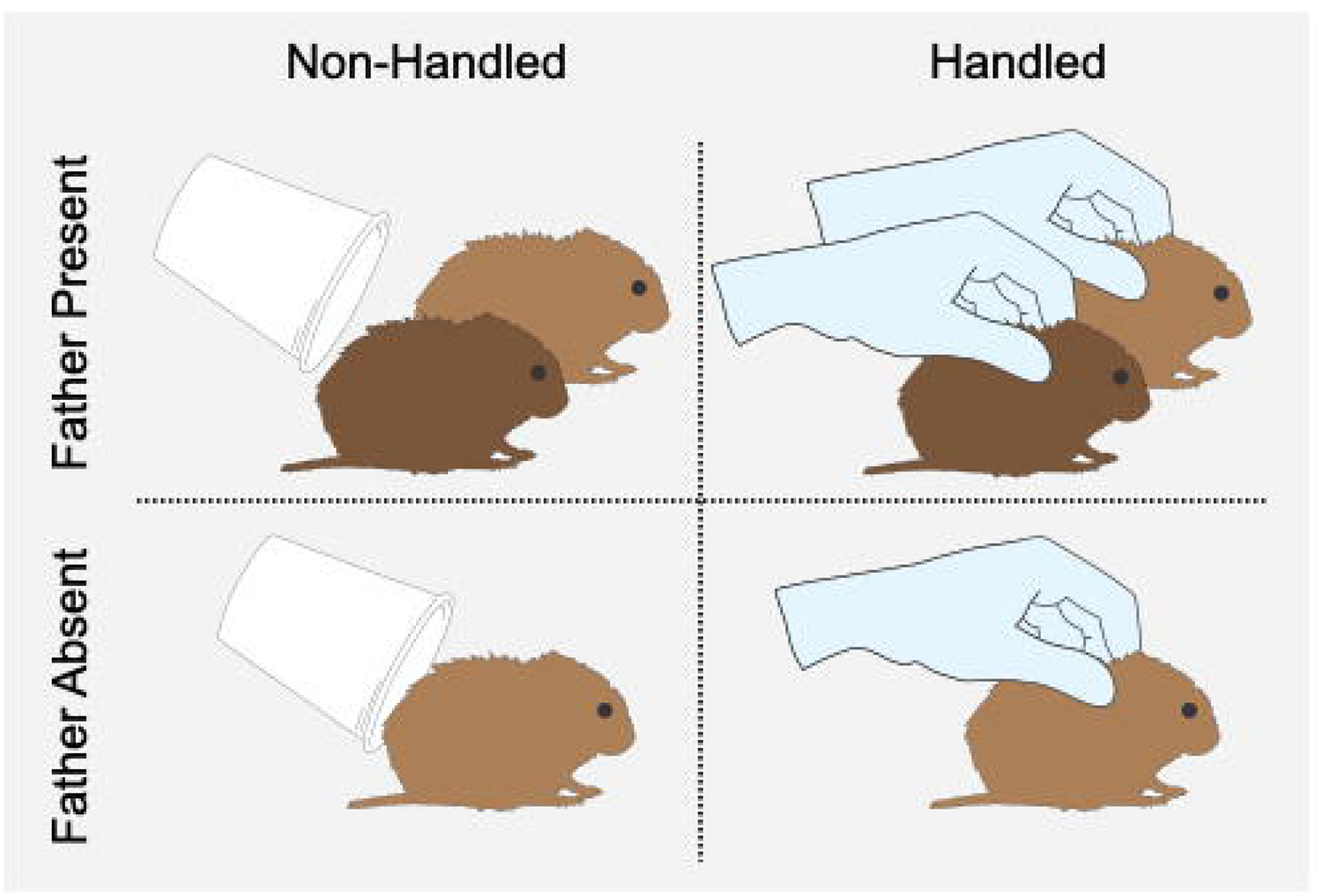
Schematic representation of the four experimental conditions. Mothers reared young in either the presence (top row) or in the absence (bottom row) of a male co-parent. During weekly cage changes, families were transferred to a clean cage either using a cup (left column) or by gently scruffing by gloved hand (right column).

### 2.3 Home Cage Analyses

One-hour home cage videos were recorded on PND2, PND9, and PND16 immediately following each cage change. We selected these postnatal days for observation because they map onto behaviorally relevant developmental timepoints (Hiura et al., 2018; Kelly et al., 2018). Briefly, PND2 pups are relatively immobile with closed eyes and are entirely dependent on parental care for food and warmth. At PND9, pups open their eyes and become more physically and socially exploratory. By PND16, pups are capable of consuming solid foods and are able to survive independently. GoPro HERO3 video cameras (GoPro Inc, San Mateo, California, USA) filmed overhead views of each family. Technical issues with the cameras resulted in several shortened videos, and thus the first 50 minutes of each video was used for subsequent behavioral analyses. Raters blind to the experimental conditions scored the behaviors of parents using Observer XT v14 (Noldus Information Technology, Leesburg, VA, USA).

### 2.4 Open-field Test

The day after pups were weaned, parents were run through a 10-minute open-field test (OFT). All tests were recorded overhead on video cameras (Sony HDR-CX330 camcorder, Sony, New York City, NY, USA). At the beginning of each test, the subject parent was gently placed by cup into the center of the transparent Plexiglas arena (57 cm x 57 cm). The arena floor consisted of a 4x4 grid of squares (14.25 cm x 14.25 cm), and the centermost four squares were designated as the arena “center” region. Video recordings were scored using EthoVision XT v13 (Noldus Information Technology, Leesburg, VA, USA) for number of visits to the center and the total distance moved in the arena.

### 2.5 Histology and Immunocytochemistry

Immediately following the OFT, parents were rapidly anesthetized by isoflurane and perfused with 0.1M phosphate-buffered saline (PBS, pH = 7.4) followed by 4% paraformaldehyde in PBS. Brains were post-fixed in 4% paraformaldehyde (24 h) then 30% sucrose (48 h) and stored at -80°C. Brains were coronally cryosectioned into three series of 40 µm slices and stored at -80°C in cryoprotectant. A single series of free-floating sections from each subject was fluorescently labeled for VP immunoreactivity (-ir). Sections were rinsed twice in PBS (30m) and blocked (1h, PBS + 10% normal donkey serum + 0.03% Triton-X-100) before being incubated in primary antibodies (48 h, Guinea Pig anti-VP 1:1000, Peninsula Laboratories, San Carlos, CA). Sections were rinsed in PBS (2 x 30m), incubated in biotinylated donkey anti-Guinea pig (1h, 1:8000, Jackson Immunoresearch, West Grove, PA), rinsed in PBS (2 x 15m), incubated at room temp in secondary antibodies (2h, streptavidin conjugated to Alexa Fluor 488 3:1000, ThermoFisher Scientific, Waltham, MA), and washed in PBS (overnight at 4°C). Sections were mounted onto microscope slides and cover-slipped with Prolong Gold antifade + DAPI nuclear stain (ThermoFisher Scientific, Waltham, MA).

### 2.6 Microscopy and quantification

Photomicrographs of the PVN and SON were taken at 10x on a Zeiss AxioImager II scope with an AxioCam MRm attachment, z-drive, and Apotome optical dissector (Carl Zeiss Inc., Gottingen, Germany). Two sections (coronal separation of 240 µm) were monochromatically imaged and manually counted for VP-ir cells using GNU Image Manipulation Program (GIMP, 2.8.22) and ImageJ (National Institutes of Health, Bethesda, MD). Cell counts of each region were combined across rostral and caudal sections of each subject and total counts per region were statistically analyzed.

### 2.7 Statistical Analysis

All analyses were conducted using R software v.4.2.2 (R Core Team, 2013). Duration data from home cage recordings and the OFT were assessed using linear mixed models (LMM) via the ‘lme4’ package (Bates et al., 2015), and p-values were derived from a likelihood ratio test within the ‘lmerTest’ package (Kuznetsova et al., 2017). Count data from home cage recordings and the OFT were analyzed using generalized linear mixed models (GLMM) assuming a negative binomial distribution with the glmmTMB package (Brooks et al., 2017), followed by type II Wald chi-squared test for significant factors. Models comparing behaviors of mothers included Father-Present/Absent condition, Experimental Handling/Non-handling condition, and postnatal day as fixed factors, and animal ID as a random factor. Models comparing behaviors between mothers and fathers included Experimental Handling/Non-handling condition, parent, and postnatal day as fixed factors, and animal ID as a random factor. Models comparing behaviors between fathers included Experimental Handling/Non-handling condition and postnatal day as fixed factors, and animal ID as a random factor. Pup grooming and pup retrievals were included as covariates in separate GLMMs with negative binomial distributions to predict cell counts, and fixed effects included handling condition (for maternal and paternal data) and father condition (for maternal data). Regression diagnostic plots and tests for over/underdispersion, heteroscedasticity, and zero-inflation were used to assess model fits, and model residuals were checked using the ‘DHARMa’ package (Hartig, 2019). Tukey corrected post-hoc contrasts and corresponding p-values of factors for both LMMs and GLMMs were extracted from the ‘emmeans’ package (Lenth et al., 2020). For all statistical models, random effects were excluded in instances where models failed to converge with their inclusion. A 0.05 α-level cutoff was used to determine statistical significance. In cases where model interactions were significant, we report the highest order interaction and related post-hoc contrast(s) in lieu of lower order effects when they involve the same factors. Three-way interactions were omitted from all models as model overcomplexity yielded convergence difficulties and risked overfitting the data.

## 3 Results

### 3.1 Total parental care pups receive from parents differs by pup age and the presence of a father

We first compared pup grooming and pup retrievals because these behaviors are exhibited by both mothers and fathers. We found that the total pup grooming a litter received from parents was dependent upon an interaction between father presence and pup age (χ2 = 6.48, df = 2, p = 0.04). Pups with fathers present received more grooming than pups without fathers at PND2 (t(97.4) = 4.55, p < 0.0001) and PND9 (t(97.4) = 2.85, p = 0.005), but not PND16 (t(97.4) = 1.54, p =0.13, **Figure 2A**).

**Figure 2.**
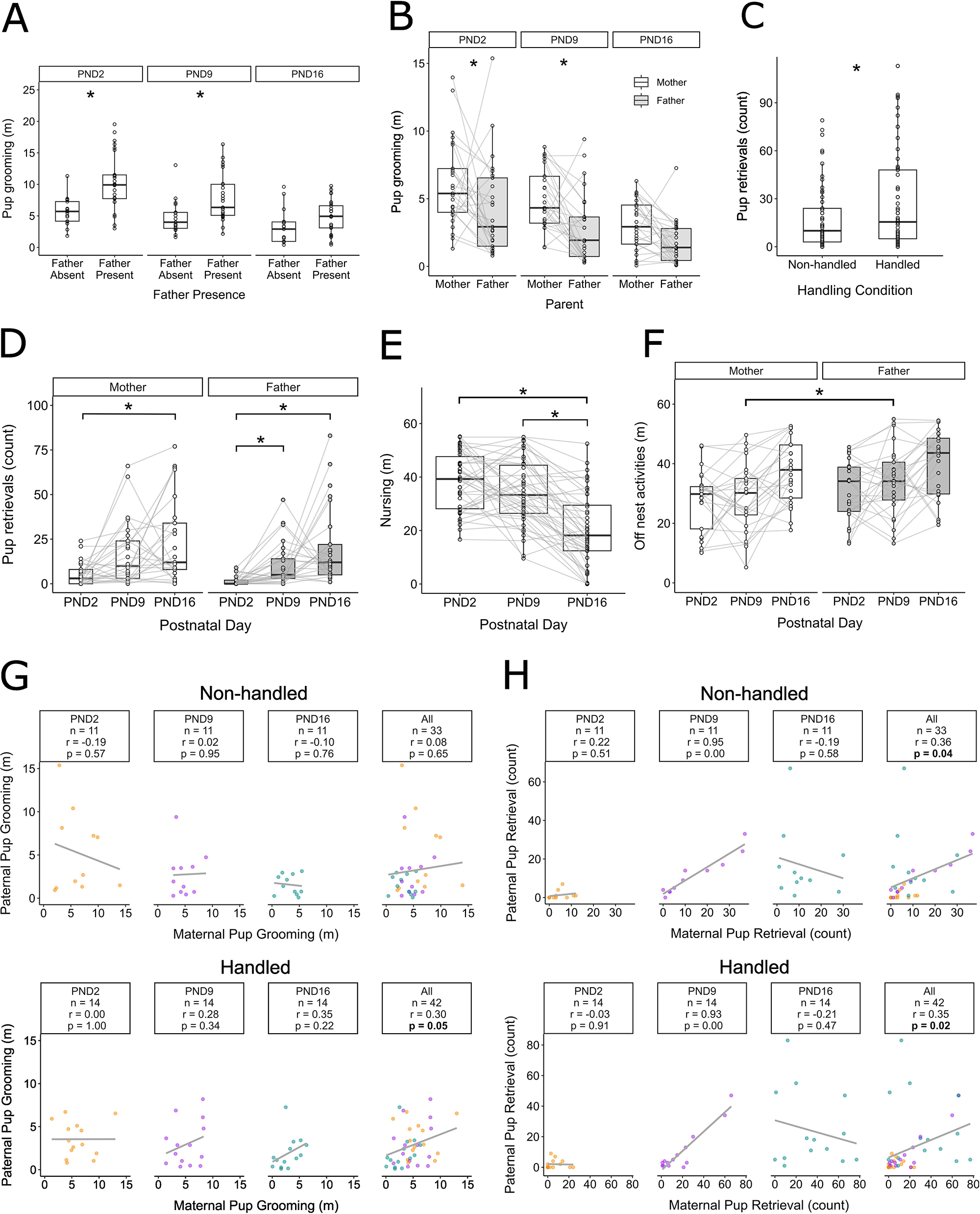
Behavioral diversity in prairie vole mothers and fathers as a function of variation in parenting experiences. (A) Total time (in minutes, m) parents groomed pups as a function of Father-Presence/Absence. (B) Total time (m) mothers vs. fathers spent pup grooming at each of the three developmental timepoints (PND2/PND9/PND16). Gray lines correspond to individual parenting pairs. (C) Counts of pup retrievals exhibited by Handled. Vs. Non-handled parents, averaging over pup age and parent sex. (D) Total maternal and paternal pup retrievals over pup postnatal days. Gray lines correspond to individual mothers across timepoints. (E) Total duration of nursing (m) by mothers over pup postnatal days. Gray lines correspond to individual mothers across timepoints. (F) Duration (m) of off-nest activity (cage exploration + trail-building) by mothers vs. fathers across pup postnatal days. Gray lines correspond to individual parents across timepoints. (G) Correlations between maternal and paternal pup grooming across postnatal days for Non-handled (top) and Handled (bottom) parents in biparental families. Yellow: PND2, Purple: PND9, Green: PND16. (H) Correlations between maternal and paternal pup retrievals across postnatal days for Non-handled (top) and Handled (bottom) parents in biparental families. *p ≤ 0.05 for all figures. Box and whisker plots and individual points visualize data distributions.

For biparental families, pup grooming differed between parents (χ2 = 18.5, df = 1, p = 1.7e-07), where mothers groomed pups significantly more than fathers did (t(117) = 4.2, p < 0.001, **Figure 2B**). Further exploratory analysis revealed that this parental difference was significant at PND2 (Mother > Father t(117) = 2.5, p = 0.01) and PND9 (t(117) = 2.98, p = 0.004), but was no longer significantly different by PND16 (t(117) = 1.86, p = 0.07). Although the number of pup retrievals did not vary by father presence (χ2= 1.82, df = 1, p = 0.18), it did differ by handling condition (χ2 = 6.2, df = 1, p = 0.01) and pup age (χ2 = 41.3, df = 2, p = 1.1e-09). Specifically, Handled parents retrieved pups more frequently than Non-handled parents (t(111) = -2.46, p = 0.02, **Figure 2C**). Total pup retrievals increased from PND2 to PND9 (t(111) = 4.63, p < 0.0001) and from PND2 to PND16 (t(111) = 6.15, p < 0.0001), but not from PND9 to PND16 (t(111) = 1.55, p = 0.37). When analyzing pup retrievals in biparental families, there was a main effect of postnatal day (χ2 = 32.6, df = 2, p = 8.3e-08) and parent (χ2 = 4.66, df = 1, p = 0.03), and a non-significant trending interaction between postnatal day and parent (χ2 = 5.22, df = 2, p = 0.07). Post-hoc tests revealed that as pups grew older, fathers generally continued to increase their pup retrievals (PND2 < PND9:t(135) = 3.6, p = 0.001), PND2<PND16: (t(135) = 5.4, p < 0.0001, PND9<PND16: (t(135) = 2.2, p = 0.07). Mothers, however, significantly differed in their pup retrievals between the ages of PND2 and PND16 (t(135) = 3.1, p < 0.008, **Figure 2D**).

### 3.2 Pup age and experiment handling, but not paternal presence, altered behavior in mothers

When comparing mothers, the total duration of nursing did not differ by paternal presence (χ2 = 0.47, df = 1, p = 0.49) or by handling condition (χ2 = 0.06, df = 1, p = 0.81). There was a significant main effect of postnatal day (χ2 = 83.2, df = 2, p < 2e-16), where PND16 pups received less nursing than they did at either PND2 (t(78) = 8.7, p < 0.001) or PND 9 (t(78) = 6.55, p < 0.001, **Figure 2E**). Maternal pup retrievals also differed between handling conditions (χ2 = 7.52, df = 1, p = 0.006), where Handled mothers retrieved their pups more frequently than Non-Handled mothers (t(111) = 2.68, p = 0.009). Fathers, on the other hand, did not alter their pup retrievals as a function of Handling (χ2 = 0.56, df = 1, p = 0.46). Additionally, the presence of fathers did not alter the amount of pup retrievals exhibited by mothers (χ2 = 1.32, df = 1, p = 0.25).

Autogrooming varied across postnatal ages for both mothers (χ2 = 13.5, df = 2, p = 0.001) and fathers (χ2 = 6.74, df = 2, p = 0.03). Mothers significantly decreased their autogrooming when pups were PND16 compared to both PND2 (t(114) = 3.6, p = 0.002) and PND9(t(114) = 3.3, p =0.004). Fathers, on the other hand, increased their autogrooming between PND9 and PND16 (t(67) = 2.4, p = 0.05). Parents also differed in the amount of time they spent exploring and trail building in the home cage (summed as “off nest” activity, χ2 = 6.0, df = 1, p = 0.01). Fathers spent more time engaging in off nest behaviors compared to mothers (t(117) = 2.6, p = 0.01), specifically at PND 9 (t(117) = 2.1, p = 0.04, **Figure 2F**).

### 3.3 Handling alters correlations between maternal and paternal pup grooming

When analyzing the relationship between maternal and paternal care combined over all pup ages, parents that were Non-handled showed no correlation in pup grooming (r = 0.08, n = 33, p = 0.65). Conversely, pup grooming was correlated for Handled mothers and fathers (r = 0.36, n = 33, p = 0.04, **Figure 2G**). Mothers and Fathers showed moderate correlations in pup retrieval when analyzing across all pup ages combined, regardless of if they were Non-Handled (r = 0.36, n = 33, p = 0.04) or Handled (r = 0.35, n = 42, p = 0.02, **Figure 2H**).

### 3.4 Handling manipulation and the presence of the father affects mothers’ behavior in the OFT

The open field test (OFT) is commonly used to assess anxiety-like behaviors, and/or exploratory behaviors under laboratory conditions. We investigated the total distance traveled within an OFT arena, and the frequency of visits to the center of the OFT arena to assess how experimental handling and the presence of fathers in the home cage impacted anxiety-like behavior in parents. The number of times mothers visited the center of the OFT chamber differed by father presence in the natal nest (χ2 = 8.5, df = 1, p = 0.004), where mothers that were co-housed with fathers visited the OFT center significantly more frequently than did single mothers (t(32) = 3.03, p = 0.005, **Figure 3A**). There was also a main effect of experimental handling (χ2 = 4.0, df = 1, p = 0.05), where Handled mothers (non-significantly) tended to visit the center of the OFT apparatus more frequently than Non-handled mothers (t(32) = 1.8, p = 0.08). Conversely, experimental handling did not impact center visits among fathers (χ2 = 0.003, df = 1, p = 0.96). Total distance traveled by mothers in the OFT was also impacted by experimental handling (χ2 = 6.5, df = 1, p = 0.01), where Handled mothers traveled significantly more than Non-handled mothers (t(34) = 2.4, p = 0.02, **Figure 3B**). A similar pattern of handling on fathers was observed, but this effect was not significant (χ2 = 3.4, df = 1, p = 0.07).

**Figure 3.**
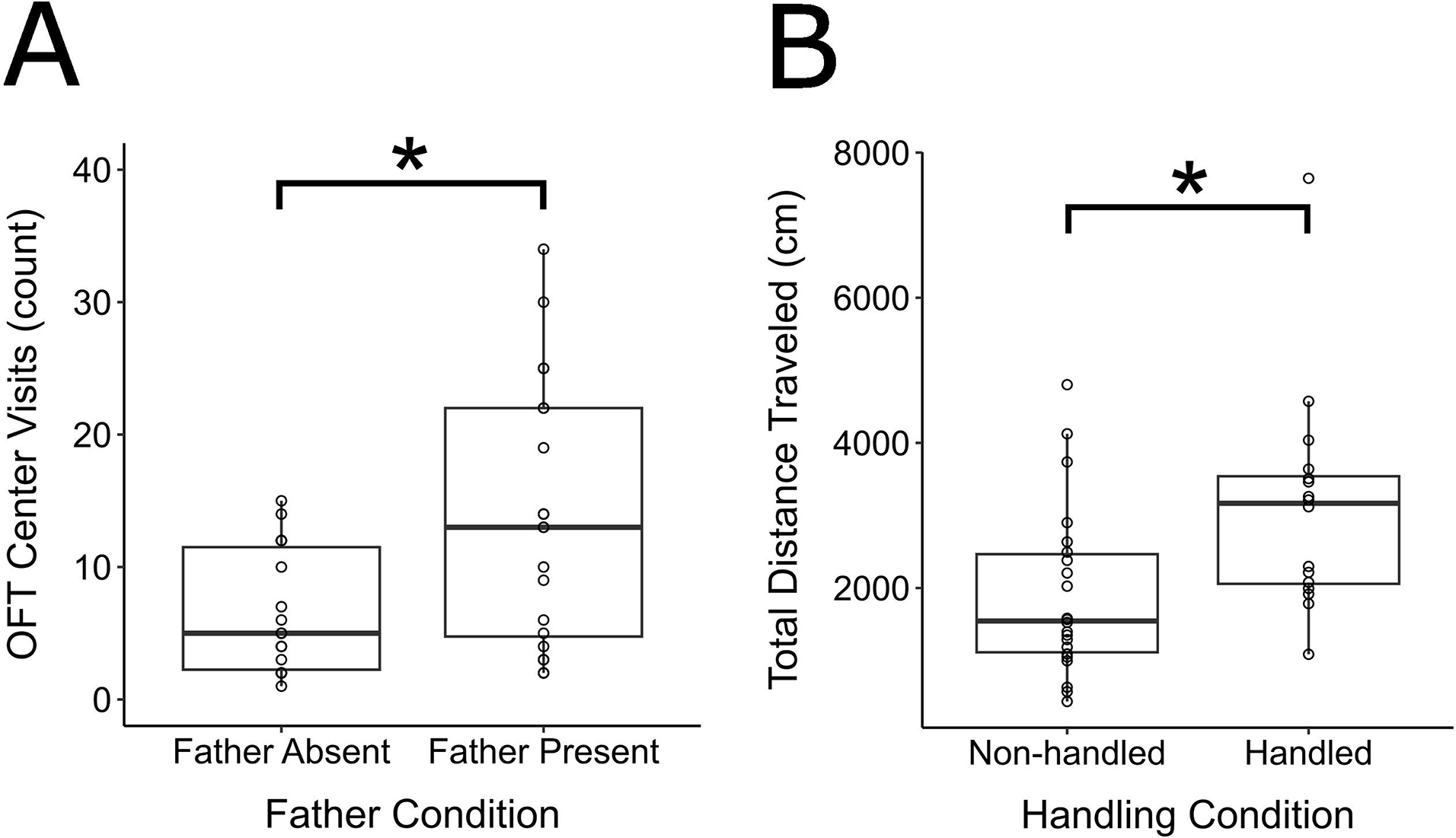
Parental Open Field Test performance across parental experiences. (A) Total number of visits to the OFT arena center by single vs. paired mothers. (B) Total distance traveled in the OFT arena by parents who were Non-handed vs. handled. *p ≤ 0.05. Box and whisker plots and individual points visualize.

### 3.5 Behavioral predictors of vasopressin cell counts varies by experimental condition

Parental care and anxiety-like behaviors have been linked to the neural activity of vasopressin systems (Bales & Saltzman, 2016; Hostetler & Ryabinin, 2013). We quantified the number of VP-ir cells in hypothalamic subpopulations to ask how their neural expression related to prior parental experiences and behaviors. In mothers, the amount of pup grooming was a significant negative predictor of the number of VP-ir cells in the PVN (χ2 = 4.0, df = 1, p = 0.05, **Figure 4A**). This relationship was independent of the presence of a partner (χ2 = 1.5, df = 1, p = 0.22) or of handling condition (χ2 = 0.8, df = 1, p = 0.37). There was no main effect of pup retrieval on maternal VP-ir counts in the PVN (χ2 = 1.1, df = 1, p = 0.29).

**Figure 4.**
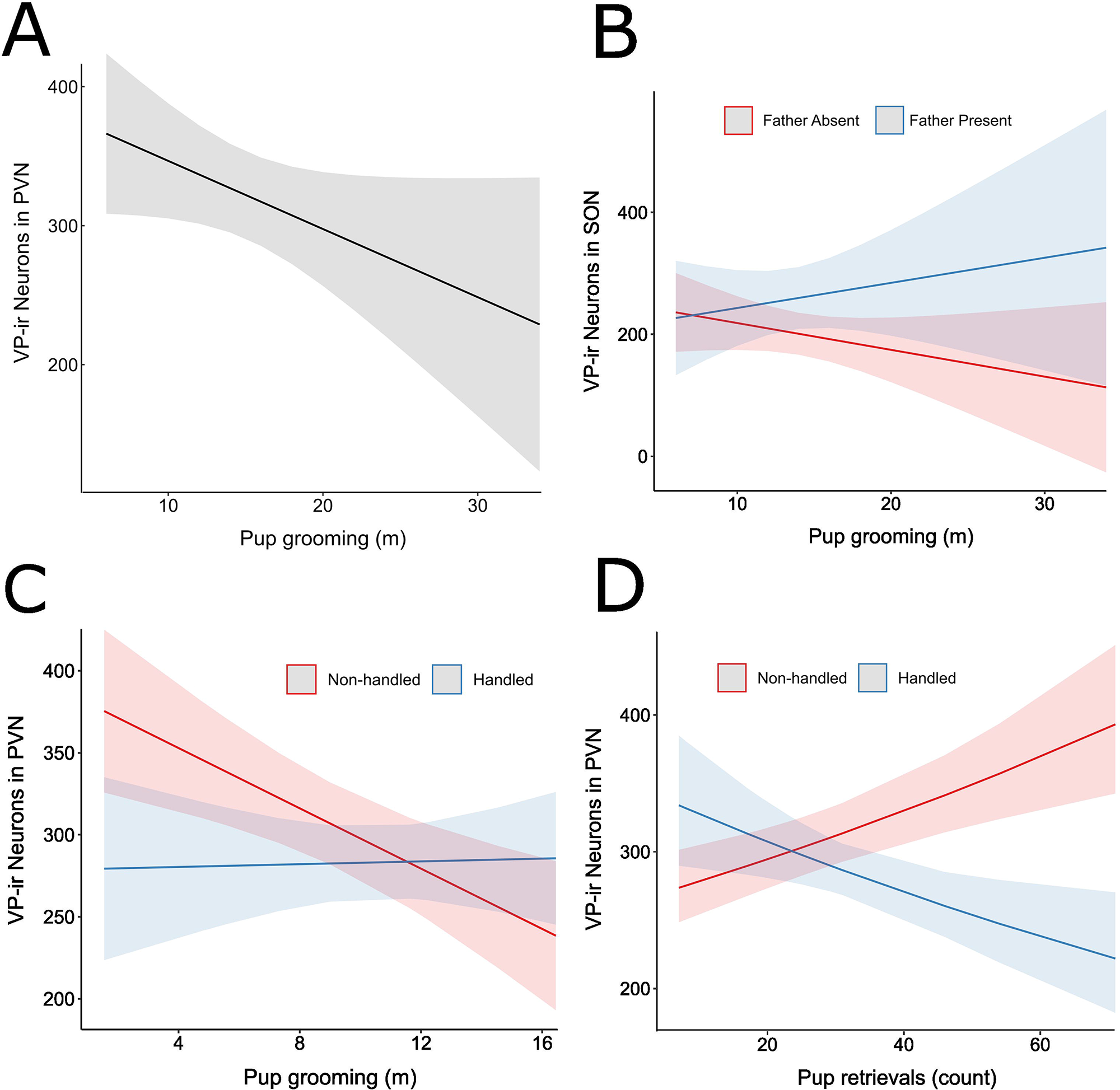
Regression visualizations for VP-ir cell counts using parental care predictors. (A) Estimated marginal means for VP-ir counts in mothers by time spent pup grooming, averaged over levels of father presence and handling condition. (B) Conditional effects on VP-ir counts by time spent pup grooming for single mothers vs paired mothers, averaged over handling conditions. (C) Conditional effects on VP-ir counts by time spent pup grooming for Handled vs Non-handled fathers. (D) Conditional effects on VP-ir counts by the number of pup retrievals by Handled vs Non-handled fathers. Shaded regions reflect 95% confidence intervals.

The direction of the relationship between pup grooming and the number of SON VP-ir cells in mothers depended upon father presence (χ2 = 4.8, df = 1, p = 0.03, **Figure 4B**). Single mothers exhibited a negative relationship between pup grooming and SON VP-ir counts, whereas paired mothers showed a positive relationship. Conversely, VP-ir counts in the SON were not related to pup retrievals in mothers (χ2 = .01, df = 1, p = 0.73)

In the fathers, the relationship between pup grooming and the number of VP-ir cells in the PVN was dependent upon the Handling condition (χ2 = 5.7, df = 1, p = 0.02, **Figure 4C**). Paternal pup grooming negatively corresponded with PVN VP-ir neurons when fathers were Non-handled, but this relationship diminished when fathers were handled. The number of SON VP-ir cells in fathers was also subject to an interaction between handling condition and pup retrievals (χ2 = 17.1, df = 1, p = 3.5e-05, **Figure 4D**). Pup retrievals were positively associated with VP-ir cell counts when fathers were Non-handled, but this association was negative when fathers were Handled. Neither pup grooming (χ2 = 0.0002, df = 1, p = 0.99) nor handling condition (χ2 = 1.7, df = 1, p = 0.2) were significant predictors of SON VP-ir counts in fathers.

## 4 Discussion

A rich understanding of the complex dynamics between parents offers insight into how interactions among caregivers alters the quality of social relationships between mating partners. Furthermore, focus on these dynamics provides additional context for how early life experiences impact offspring social behavior and wellbeing. The present study examined 1) how variation in (co-)parenting experiences for mothers and fathers impact their behaviors inside and outside of the natal nest, and 2) how parental experiences shape the neural systems known to be important for parental behavior and anxiety-like traits.

### 4.1 Parents exhibit offspring age-dependent behavioral plasticity in sex-specific ways

We used a 2x2 design integrating the presence and absence of fathers with a salient but minor handling stress manipulation designed to temporarily alter parental behavior (Bales et al., 2007) to model ecologically relevant and distinct rearing conditions (Getz & Carter, 1996). We also investigated the changes in caregiving under these conditions over the course of offspring development. Despite these experimentally imposed conditions, offspring age was the factor that most frequently accounted for significant variation in parental and non-parental behaviors. These results are consistent with prior work showing that prairie vole parents display different rates of care when their pups are neonates compared to when they are older (Lonstein & De Vries, 1999). This phenomenon is not unique to biparental voles; many other species have demonstrated robust parental behavioral plasticity in response to offspring age. For example, zebra finch parents whose chicks were swapped with broods that were either older or younger than their own offspring adjusted the duration of their caregiving period to match the needs of their foster brood (Rehling et al., 2012). Results such as these support the notion that the capacity to respond to the changing needs of the offspring can serve an adaptive function for species that exhibit parental care (Saltzman et al. 2017). There are several ways in which parents can balance the energetic demands of rearing pups with their need to maintain their own body condition and maximize future reproductive potential (Trivers, 1974). For example, this can be accomplished by investing effort to meet the current needs of offspring while decreasing behaviors that no longer appreciably support them. In line with this balancing act, we found that parents decreased caregiving behaviors, including nursing and grooming, as the pups approached an age associated with the ability to self-thermoregulate and consume solid food. Conversely, pup retrieval by both parents increased as pups grew older, a result likely attributable to the refinement of pup motor skills and the onset of their exploratory behaviors. The capacity to decrease the time allocated to intense offspring-directed care might also explain the observed increases in off-nest activities – a trade-off that has been theoretically modeled and empirically tested in other species (Kacelnik & Cuthill, 1990; Whittingham, 1993; Williams, 1966).

Thomas and Birney (1979) argued that male and female prairie voles contribute many of the same behaviors to offspring care. Other experiments have since shown that, compared to mothers, fathers tend to be more variable in the degree to which they care for offspring (e.g., Solomon, 1993; Kelly et al. 2020; Finton & Ophir 2020). Consistent with all of this, we report sex differences in the patterns of adjustments in pup age-related behaviors, indicating that mothers and fathers exhibit differential sensitivities and responses to offspring developmental stage. Specifically, mothers groomed pups more than fathers did over the course of development, particularly when pups were young (2 days old and 9 days old). This is in line with earlier work demonstrating that prairie vole mothers spend more time in contact with pups than fathers do during the first week of postnatal life, but not beyond (Solomon, 1993). We also found that fathers increase their autogrooming as pups age, whereas mothers decrease their autogrooming over the same timeframe. Notably, a contrast appears to exist between mothers that prioritize offspring care (i.e., offspring grooming) over self-care (i.e., autogrooming), and fathers that more readily shifted from pup care to self-care behavior. This difference between parents suggests that despite engaging in the same general behaviors, fathers of this socially monogamous and biparental species invest relatively less in their offspring and relatively more in themselves compared to mothers. Many of the mechanisms underlying the expression and diversity of parental behavior appear to be sex-specific (Bendesky et al., 2017). Presumably distinct selective pressures on neural mechanisms such as these may have shaped how parents respond to pups of different ages (i.e., age-dependent behavioral plasticity) (Trivers, 1974). Whatever the reason, our data indicate that parental sex differences exist and can be tracked to when sex-specific trade-offs occur between pup-directed vs self-directed care behaviors.

### 4.2 Absence of fathers altered mothers’ anxiety-like, but not pup-directed, behaviors

For mammalian mothers who invest heavily in offspring, co-parenting partners can offset rearing costs and offspring mortality by provisioning resources, defending territories against intruders and predators, providing thermoregulatory and social stimulation, and tending to the young, thereby freeing mothers to engage in non-parental behaviors (Woodroffe & Vincent, 1994). In some mammals, paternal care enhances offspring maturation rates, litter sizes, body condition, and survival (Gubernick et al., 1993; Stockley & Hobson, 2016; Wang & Novak, 1992), often as a result of shared or added forms of care. Several models assessing the evolution of biparental care predict that single parents of biparental species should increase their caregiving behaviors to (at least partially) compensate for the absence of a mate (McNamara et al., 1999). Support for these models showing that single mothers engage in more pup directed behavior compared to partnered mothers has been found among degus (*Octodon degus*), Mongolian gerbils (*Meriones unguiculatus*), rock cavies (*Kerodon rupestris*), and the Mexican volcano mouse (*Neotomodon alstoni*) (Elwood & Broom, 1978; Luis et al., 2004; Tasse, 1986; Wilson, 1982). Notably, single prairie vole mothers did not adjust maternal care due to the absence of fathers in the current study. Indeed, we found no evidence that mothers behaviorally upregulate their care to compensate for the absence of a father, echoing the findings of several other studies of prairie vole parental care (Ahern et al., 2011; Ahern & Young, 2009; Bosch et al., 2018; McGuire et al., 2007; Rogers & Bales, 2019; Tabbaa et al., 2017). Moreover, one study on biparental California mice also found that single mothers do not differ in their maternal care compared to paired mothers (Zhao et al., 2019). These results raise the question: why did prairie vole single mothers not upregulate their caregiving to compensate for the absence of a father? One hypothesis is that the laboratory conditions under which mothers were tested resulted in a ceiling effect that lowered the costs required for sufficient offspring care. Implementing sufficiently harsh or challenging environmental conditions may therefore force single mothers to tradeoff pup care and self-care (Wright & Brown, 2002). Doing so would presumably reveal the full potential of paternal presence on both pups and maternal care, a hypothesis that is supported by recent evidence (Kelly et al., 2020). Whether our laboratory conditions were insufficient to induce changes in prairie vole maternal care or not, our data contrasted with other studies that revealed that many biparental rodents show maternal compensation for the absence of fathers in the natal nest. This phenomenon raises another unresolved question: when mothers *do* compensate for the lack of a co-parent, are they responding to altered pup behaviors as a function of paternal presence, or to the direct absence of the fathers themselves (Elwood & Broom, 1978)? Additional work focusing on dissociating the contextual features to which parents may be adapting is necessary to identify the cues that may elicit parental behavioral plasticity.

Although paternal absence did not appear to modify maternal care, paired mothers visited the center of the OFT chamber more frequently than single mothers. This behavior is often interpreted as representing a less “anxious-like” phenotype. Because single and partnered mothers did not display other differences in behavior within their home cages, it is possible that anxiety-related behaviors outside of a parental context may be more susceptible to perturbation by experience during the parenting period. These results concur with earlier work in which single mothers show greater levels of anxiety-like and depressive-like phenotypes compared to paired prairie vole mothers in an elevated plus maze and forced swim test, respectively (Bosch et al., 2018). However, our OFT testing followed pup-weaning, whereas Bosch et al. (2018) measured behavior in the days following parturition. In both experimental designs, it is challenging to discern if the behavioral results are due to bond dissolution or to the experience of single parenthood. When taken together, these studies suggest that the interval of time between partner separation and testing for anxiety-like traits may influence the direction of the effects of single-motherhood. It would be useful to address the potential differential influences of partner loss versus the demands of single-parenthood to best understand how these factors independently or synergistically shape parental anxiety-like phenotypes.

### 4.3 Acute handling stress induced coordination of pup grooming between mothers and fathers

Experimental handling significantly increased the activity of parents in terms of caregiving (exemplified by more pup retrievals in the home cage) and general activity (i.e., distance traveled in the OFT). Notably, handling produced a correlation between maternal and paternal pup grooming that was not observed when parents were not handled. This might suggest that behavioral upregulation of grooming induced by the acute handling paradigm promoted coordination of some aspects of caregiving between parents. However, parental correlations in pup retrievals emerged independently of handling experience, which suggests that if the acute stressor synchronized parental care, it did so in a behavior-specific way. Behavioral coordination or synchrony in biparental species is dependent on social experience and context (Prior, 2020). For example, experimentally manipulating brood size led to changes in the synchronization of nest visits in wild zebra finch parents (Mariette & Griffith, 2015). Furthermore, nest visit synchrony in blue tits (*Cyanistes caeruleus*) has been linked to weather and altitude, suggesting that biparental coordination is sensitive to ecological conditions (Lejeune et al. 2019). Our data (in conjunction with the aforementioned studies) supports the hypothesis that dyadic care exhibits experience-specific and context-specific plasticity. Little is known regarding the neuroendocrine mechanisms that may facilitate these adaptations, but below we explore the evidence that suggests a potential link between VP cells and experience-dependent neurobehavioral plasticity in parents.

### 4.4 Associations between vasopressin cell counts and parental behaviors exhibited experience-dependent plasticity in mothers and fathers

Our data demonstrated that both mothers and fathers exhibited predictive relationships between their home cage caregiving behaviors and hypothalamic vasopressinergic cell counts. The duration that mothers spent grooming their pups was inversely associated with the number of VP-ir neurons in the PVN, suggesting that there may be a negative relationship between vasopressin signaling and maternal pup grooming. Considering that VP is involved in parental status and behavior in prairie voles (Bamshad et al., 1994; Wang et al., 1994, 2000) and other rodents (Bayerl et al., 2016; Bosch & Neumann, 2008; Parker & Lee, 2001), it initially seems paradoxical that mothers that expressed higher levels of grooming had fewer VP-ir cells. Interpretations of cell count data are quite challenging because differences in immunoreactivity can represent either peptide production, accumulation due to blocked secretion, or changes in release (Goodson & Kabelik, 2009; Panzica et al., 2001). Under the latter interpretation, the decrease in PVN VP-ir neurons that we found could indicate greater release of bioavailable VP to extrahypothalamic sites, which in turn might have facilitated increases in maternal care. To determine the functional significance of variation in VP-ir density, future studies could utilize acute sampling methods or imaging of fluorescent reporters to determine how real-time fluctuations in VP release relate to the expression of distinct parental behaviors.

Like with the PVN, VP-ir counts in the SON of mothers were also significantly associated with pup grooming. In contrast, however, the direction of the association between pup grooming and VP-ir cells counts was contingent on whether mothers raised offspring with a co-parent or raised offspring alone. Similar experience-dependent interactions of cell counts and parental care were observed in fathers, in which the handling conditions determined if the slope between VP-ir counts and pup grooming/pup retrievals were positive or negative. Critically, our neural measures took place after the behavioral observation periods and are therefore only correlational, making it challenging to ascertain how the associations between neural and behavioral variables relate to VP-mediated functional control of parental care. Nevertheless, our results reveal a general pattern in which social and nonsocial experiences shaped the associations between parental behavior and hypothalamic VP cell groups. Similar patterns of experience-dependent mediation of brain-behavior relationships have been described for several neuromodulators in other species. For example, a study of wire-tailed manakins (*Pipra filicauda*) found that the relationship between circulating testosterone and male social behaviors was inverted between territory-holders and non territory-holders, indicating that social status dynamically modulates hormone-behavior relationships in this species (Ryder et al., 2020). Furthermore, the association between baseline corticosterone levels and parental success (fledgling numbers) is positive prior to egg laying and negative during the subsequent parental provisioning phase in great tits (*Parus major*), and is therefore dependent on reproductive stage (Ouyang et al., 2013). Moreover, group-housed mice show a significant positive correlation between serotonergic activity and social investigation, but such a correlation was not found in isolate-housed mice, suggesting that social housing contexts coordinate relationships between serotonin and social behavior (Keesom et al., 2017). Our study supplements these examples supporting the hypothesis that social and stress-related experiences are involved in contingently shaping the relationships between neuropeptides and behavioral phenotypes, even across neurotransmitter classes and taxa.

The ways in which our manipulations of parental experience mediated the direction of brain-behavior phenotypes was different between fathers and mothers. How experience-dependent plasticity manifests in the brains of parents may be tied to sexual dimorphisms in neural and behavioral phenotypes. The majority of what is known about the neuroendocrinological basis of mammalian parental behavior has been characterized in females (Saltzman & Ziegler, 2014). From these studies, it is clear that a symphony of hormones and neuro active signaling molecules accompany pregnancy and parturition, and are critical for priming primiparous mothers to behave maternally toward their newborn offspring. Mammalian fathers, on the other hand, do not gestate their young. Thus, the endocrine mechanisms that subserve care in males can greatly differ from those of females, and might vary between species (Horrell et al., 2018). Nevertheless, hormones known to play a functional role in the display of paternal behaviors do fluctuate and are susceptible to factors such as mating and pair bonding experiences, exposure to pregnant females, and exposure to offspring (Saltzman & Ziegler, 2014). The fact that the VP-behavior relationships we observed were contingent upon other environmental experiences (presence of a co-parent in mothers, and handling stress in fathers) highlights the remarkable plasticity of the vasopressin signaling system, even within the brains of mature animals. It is noteworthy that the highly variable distributions of forebrain vasopressin receptors across species stands in stark contrast to the deeply conserved patterning of other neuromodulatory systems such as steroid hormone and dopamine receptors (King & Young, 2016). This evidence in conjunction with our data lends credence to the notion that VP systems exhibit high levels of evolvability that may enable this hormonal signal to take on new functions in the modulation of social behavior (Young & Zhang, 2021).

Taken together, we investigated how social and non-social variation in parental experiences impact prairie vole parents’ behaviors across pup development, and how these experiences shape the hormonal systems that drive and respond to social behaviors. Although manipulating parental composition and acute stress did not modulate parental behavior as strongly as pup age did, these variables robustly shaped the associations between VP cell density and pup-directed care in mothers and fathers. Our findings extend what is known about the interconnections between parental contexts, behaviors of parents, and neuroendocrine phenotypes, and demonstrates the value of integrating behavioral and physiological measures to understand the longitudinal dynamics of brain-behavior relationships. Future work should address the functional implications of alterations in hypothalamic neuroendocrine signaling for parental behavior, and search for the underlying mechanisms by which experiences biologically organize the relationships between hormones and parental behavioral phenotypes. Comparisons of interclass species may also reveal to what degree such evolutionary mechanisms may be shared across biparental species.

## 5 Author contributions

LCH and AGO conceived, designed the experiments, and discussed the results. LCH conducted behavior testing, brain sectioning, histology, and data analysis. VAL assisted with behavioral scoring and brain sectioning. LCH and AGO interpreted the results and wrote the manuscript. All authors critically revised the article and approved the final version.

## 6 Funding

This work was supported by funding from the Eunice Kennedy Shriver National Institute of Child Health and Human Development to AGO (HD079573) and from an NSF Graduate Research Fellowship to LCH (1650441).

## 7 Conflict of interest

The authors declare no conflict of interest.

## Acknowledgements

We would like to thank Mandy Chan for her assistance with behavioral scoring. Thank you to the animal care staff of Cornell University for making this research possible, and to the voles for their critical role in advancing scientific knowledge.

## Notes

### Competing Interest Statement

The authors have declared no competing interest.

